# Comprehensive investigation of AAV tropism across human iPSC-derived neuronal subtypes

**DOI:** 10.64898/2026.03.24.713895

**Authors:** Linus Wiora, Salvador Rodriguez-Nieto, Leelja Rößler, Jacob Helm, Alejandra Leyva, Thomas Gasser, Ludger Schöls, Ashutosh Dhingra, Stefan Hauser

## Abstract

Recombinant Adeno-associated viruses (AAVs) are widely used for gene delivery in the central nervous system and have become central tools in both gene therapy and basic neuroscience research. However, although AAV serotypes have been extensively characterized in rodent models, their performance in human neurons, particularly those derived from induced pluripotent stem cells (iPSCs), remains poorly characterized. While human iPSC-derived neurons are increasingly used for disease modeling and drug screening, their susceptibility to viral transduction varies and remains difficult to predict.

In this study, we systematically evaluated the transduction efficiency and toxicity profiles of 18 wild-type and engineered AAV serotypes across three distinct types of iPSC-derived neurons, relevant to disease modeling and drug discovery: cortical projection neurons, NGN2- induced forebrain-like neurons, and dopaminergic neurons and four doses (1E3, 1E4, 1E5 and 2E5 genome copies per cell). Using automated high-throughput confocal imaging and quantification of reporter gene expression, we identified several serotypes with robust and efficient transduction across all neuronal subtypes. Among these, three serotypes AAV6, AAV6.2 and AAV2.7m8 showed consistently high performance.

To assess safety, we quantified cell number and neurite morphology, finding that while high transduction and gene expression correlate with toxicity, sensitivity varied across neuronal subtypes, with NGN2 neurons being most vulnerable and dopaminergic neurons most resilient. Finally, we validated our findings in a more complex 3D model by testing one of the best-performing serotypes, AAV2.7m8, in both whole and dissociated human cerebellar organoids.

Together, our results establish a benchmark dataset for AAV performance in human iPSC-derived neurons and provide practical guidance for AAV based gene delivery in human *in vitro* neural models. This resource will be valuable for both basic research and preclinical applications aiming to manipulate gene expression in human neurons and understanding AAV tropism in disease-relevant cell types.

## Introduction

Adeno-associated viruses (AAVs) are small, non-enveloped, replication-defective particles belonging to the genus *Dependovirus*. They package a ∼4.7 kbp single-stranded DNA genome, and wild-type variants can be found in humans and other primates (Salganik et al., 2015; Sant’Anna & Araujo, 2022). Due to their broad tropism across human tissues and favorable safety profile, AAVs have gained widespread use in gene therapy applications (Au et al., 2022; Naso et al., 2017), including numerous clinical trials and several approved treatments for genetic diseases (Wang et al., 2024).

Beyond therapeutic applications, AAVs are widely used as vectors to deliver genetic tools in the central nervous system (CNS) of animal models (Challis et al., 2022; Haggerty et al., 2020). They have enabled precise control of neuronal activity via optogenetics and are valuable for circuit mapping due to the neuronal specificity and retrograde transport of certain serotypes (Haery et al., 2019; Minetti, 2024). However, findings from rodent studies often fail to fully translate to primates and/or humans (Drouyer et al., 2024; Fang et al., 2025). The advent of induced pluripotent stem cells (iPSCs) - which can be reprogrammed from somatic cells and differentiated into nearly any cell type - offers new opportunities to study human biology and disease in a genetically relevant and cell type specific context (Okita et al., 2011; Takahashi et al., 2007).

For neurological disorders, patient-derived iPSC neurons enable *in vitro* disease modeling of affected cell types. Several protocols have been developed to generate neurons from iPSCs, which broadly fall into two categories: (1) growth factor–based protocols that mimic developmental signaling to induce specific neuronal fates, and (2) transcription factor-based protocols that overexpress key regulators of neuronal differentiation. While the former can yield well-defined neuronal subtypes, they are typically lower in throughput. In contrast, overexpression-based approaches produce neurons more rapidly and at higher yield, making them suitable for screening applications (Flitsch et al., 2020), though the resulting populations may be more heterogeneous (Lin et al., 2021).

While protocols for delivering genetic material to undifferentiated iPSCs are well established, introducing transgenes into terminally differentiated neurons remains a significant challenge. Mature neurons are highly sensitive to conventional transfection methods such as electroporation or lipofection, often resulting in poor viability and low transduction efficiency (Alabdullah et al., 2019). Although creating stable, inducible iPSC lines can circumvent these issues, this approach is labor-, time-, and cost-intensive.

AAVs are a promising alternative for gene delivery to iPSC-derived neurons. However, limited success in standard *in vitro* models has led to their underuse in this context. While some studies have reported AAV-mediated transduction of iPSC-derived neurons (Fischer et al., 2022; Weinmann et al., 2022), a systematic comparison of common AAV serotypes across different neuronal differentiation paradigms and across a broad dose range, combined with quantitative toxicity readouts, remains limited.

Here, we present a comprehensive screen of 18 wild-type and engineered AAV serotypes in three distinct iPSC-derived neuronal subtypes and four doses. Our systematic approach reveals that the transduction efficiencies observed in these human neuronal cultures often deviate significantly from the expected neurotropism reported by prior *in vivo* studies. Specifically, we find that some AAVs highly regarded for their in vivo neurotropism are poorly effective in the iPSC platform, while others developed for non-neuronal tissues demonstrate surprisingly robust efficiency. These findings indicate a critical gap in our understanding of AAV tropism and specificity and highlight the value of the iPSC model for studying AAV biology in a human-relevant context. We identify serotypes with robust transduction efficiency, establishing a valuable resource for both basic neuroscience and preclinical gene therapy research. Finally, we test whether the top-performing vector retain strong transduction capability in a 3D setting using iPSC-derived cerebellar organoids. These optimized vectors hold potential for improving the genetic accessibility of human neurons i*n vitro* and may inform the selection of clinically relevant AAVs for CNS-targeted therapies.

## Results

### Characterization of iPSC-derived neuronal subtypes confirms efficient and reproducible differentiation

To establish a robust platform for AAV screening, we first differentiated human induced pluripotent stem cells (iPSCs) into three distinct neuronal subtypes commonly used in the field: cortical projection neurons (Rehbach et al., 2019; Shi, Kirwan, & Livesey, 2012; Shi, Kirwan, Smith, et al., 2012), forebrain glutamatergic neurons (Zhang et al., 2013) and dopaminergic neurons (Dhingra et al., 2020; Reinhardt et al., 2013).

Immunocytochemistry and confocal imaging confirmed the expression of subtype-specific neuronal markers. All neuronal subtypes used in this study showed positive immunostaining for the pan-neuronal marker β-III-tubulin (TUJ). Cortical neurons (CN) showed high expression of the cortical layer specific marker CTIP2. NGN2 neurons were positive for the neuronal markers TUJ and MAP2, while dopaminergic neurons (DA) stained positive for TH. Morphologically, all subtypes exhibited complex neurite networks consistent with mature neuronal phenotypes (Fig. 1A).

**Figure 1:**
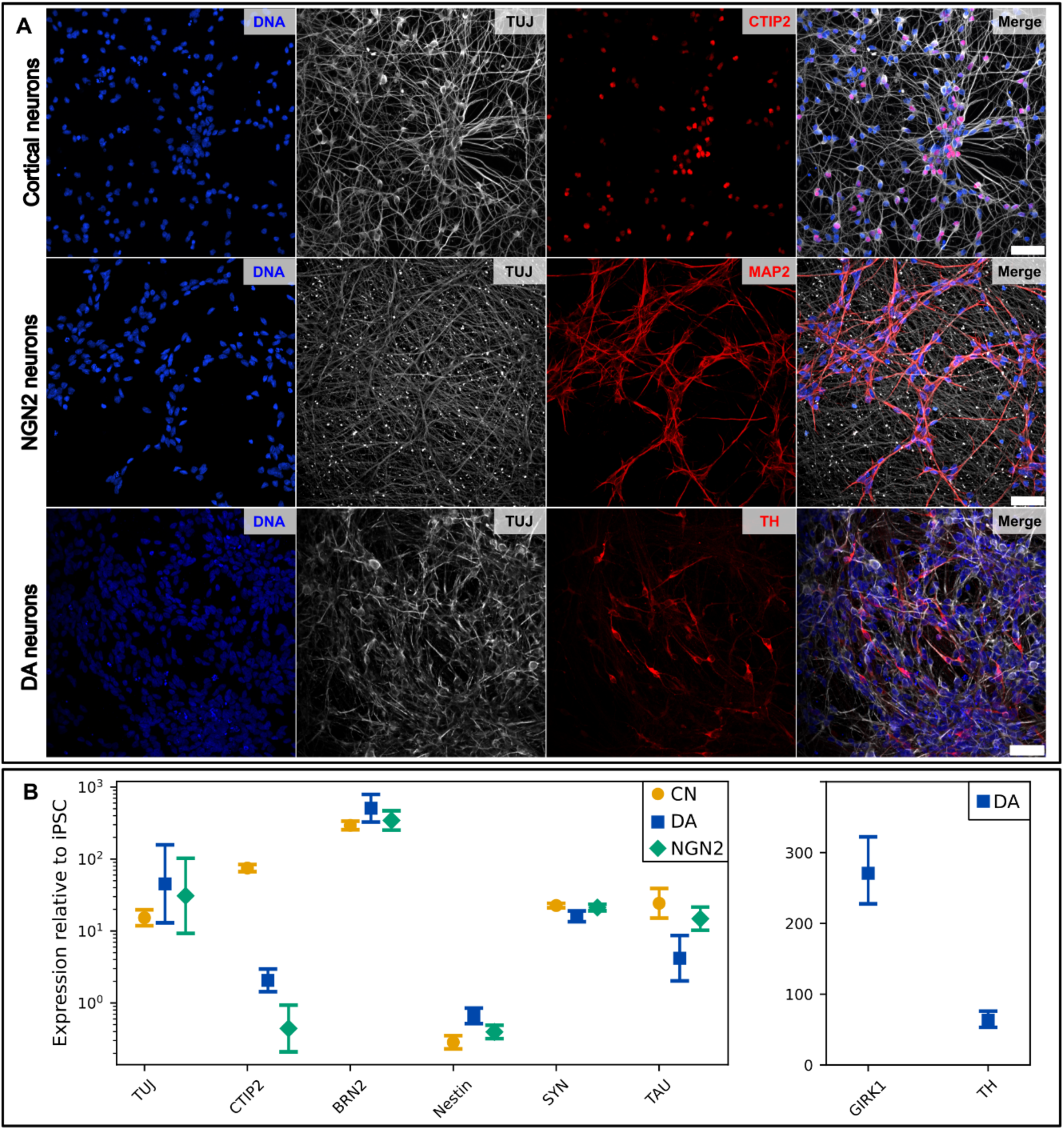
**Characterization of iPSC-derived neuronal subtypes**: cortical neurons (CN), dopaminergic neurons (DA) and NGN2 derived neurons (NGN2). **A:** Immunocytochemistry showing expression of the pan-neuronal markers TUJ/MAP2 and specific markers for cortical (CTIP2) and dopaminergic (tyrosine hydroxylase, TH) neurons (Scale bar: 50 µm) **B:** Gene expression relative to GAPDH and TBP assessed by qPCR for neuronal markers. All neuronal subtypes express general neuronal markers such as TUJ and BRN2 as well as markers of maturity like synapsin (SYN) while the expression of the neural precursor marker NESTIN remains low. Linage specific markers are expressed in the respective cell type (CTIP2 in CN, TH/GIRK1 in DA, not detected in CN and NGN2).

Gene expression analysis by qPCR revealed high mRNA levels of neuronal marker genes compared to iPSCs. Notably, *CTIP2* expression was elevated specifically in cortical neurons (∼100 fold), while GIRK1 and TH were only detectable in dopaminergic neurons, highlighting subtype-specific transcriptional profiles. Low levels of the precursor marker *NESTIN*, together with increased expression of *BRN2*, *SYNAPSIN* and *TAU*, were indicative of mature neuronal cultures (Fig. 1B).

This validated system formed the foundation for downstream AAV transduction studies. Having established a reliable platform of three well-characterized neuronal subtypes, we next performed a systematic AAV serotype screen using automated high-throughput confocal imaging to quantify transduction efficiency.

### Systematic AAV serotype screening reveals cell type-specific transduction profiles

To assess the transduction efficiency of different AAV serotypes in human neurons, we conducted a systematic screen using 18 commonly used AAV variants at four different doses (multiplicity of infection [MOI] of 1E3, 1E4, 1E5 and 2E5 genome copies [gc]/cell) across three iPSC-derived neuronal subtypes (Fig. 2A). All AAV serotypes contained an identical GFP expression cassette under the control of the CAG promoter, enabling a screenable readout for transduction rate via confocal microscopy. Terminally-differentiated neurons were transduced, and transgene expression was monitored by longitudinal live-cell imaging for the highest dose and endpoint confocal microscopy in the GFP channel for all dosages.

**Figure 2:**
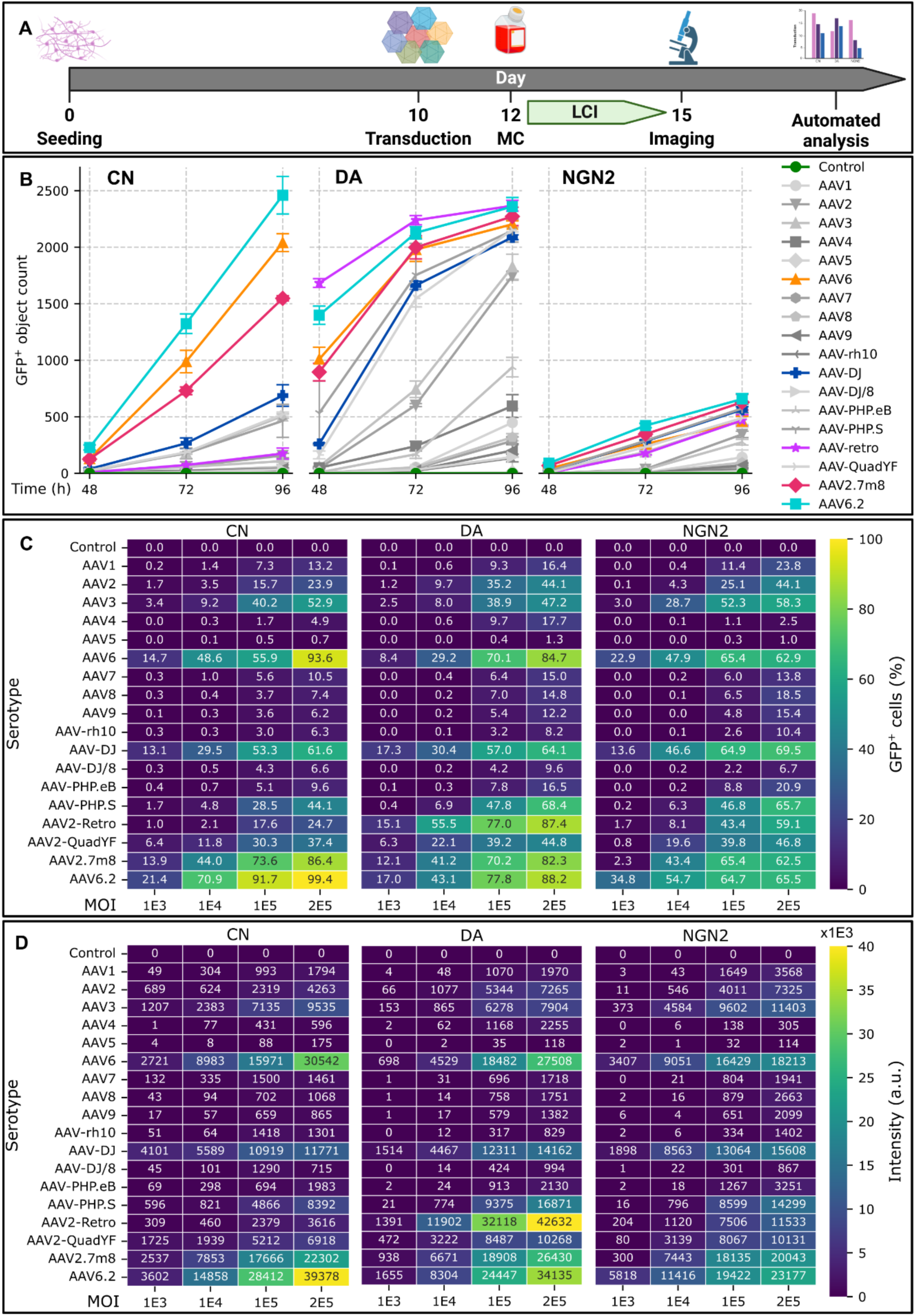
Comparison of AAV transduction across neuronal subtypes. **A:** Schematic outline of experimental procedure. Neurons were seeded and matured in 96 well plates, transduced with the AAV panel and imaged after medium change (MC) 2 days post treatment. **B:** GFP+ object count over time determined by live cell imaging starting 48h after AAV treatment for the highest AAV dose (MOI 2E5). **C** AAV transduction rate and **D:** GFP intensity at endpoint measured by confocal imaging (n=3 wells, 16 FOV/well). AAV6, 2.7m8 and 6.2 are efficiently transducing all neuronal subtypes tested. AAV2-Retro shows high specificity for dopaminergic neurons.

Live imaging revealed time-dependent increases in GFP expression following AAV transduction, with clear differences in onset kinetics and overall efficiency across serotypes (Fig. 2B). The strongest increases in fluorescence were observed with AAV6, 6.2, and 2.7m8, particularly in cortical neurons. These serotypes were outperformed only by AAV2-retro selectively in dopaminergic neurons. No significant cytotoxicity or morphological changes were observed during the time course, indicating good tolerability across all tested conditions.

To systematically compare transduction efficiency, we quantified the percentage of GFP- positive cells in confocal images acquired at the experimental endpoint 48h after AAV transduction. Heat maps summarizing the transduction rates for all 18 serotypes across four viral doses revealed distinct patterns of serotype performance patterns in each neuronal subtype (Fig. 2C). Some serotypes, such as 6.2 and 2.7m8, exhibited consistently high transduction across all cell types, while others demonstrated pronounced subtype specificity. For example, AAV2-retro showed preferential tropism for dopaminergic neurons but not for cortical or NGN2 neurons.

Systematic quantification revealed that the highest transduction efficiencies were consistently achieved by AAV6 (representing a high-performing wild-type serotype) and the engineered variants AAV6.2 and AAV2.7m8. In cortical neurons, AAV6 and AAV6.2 were the most effective vectors, achieving transduction rates of 93.6% and 99.4%, respectively, at the highest dosage, significantly outperforming the next closest serotype. In NGN2 neurons, AAV.DJ, a serotype engineered for liver specificity (Grimm et al., 2008), was highly efficient, reaching a transduction rate of 69.5%. Meanwhile, in dopaminergic neurons, AAV2-retro showed the greatest cell-type specificity, achieving a transduction rate at the highest dosage (87.4%) that was only marginally lower than the overall best performer, AAV6.2 (88.2%). Notably, at an intermediate MOI of 1E4 gc/cell, AAV2-retro even outperformed AAV6.2 in this specific neuronal subtype.

In addition to the percentage of positive cells, we also quantified the mean fluorescence intensity (Fig. 2D). As expected, the transduction rate and intensity (representing gene expression) correlated strongly for the three neuronal subtypes (*r* = 0.95, 0.96 and 0.93 for CN, DA, and NGN2, respectively). In cortical neurons, AAV6.2 that showed the highest transduction rate also exhibits the highest fluorescence intensity. However, in dopaminergic neurons, AAV6.2 transduced the most cells; yet, AAV2-retro outperformed this serotype in terms of intensity, showing the highest fluorescence intensity, and therefore, gene expression of all conditions tested (MOI 2E5). NGN2 neurons generally showed lower intensity values in direct comparison, reflecting the lower transduction rates achieved with this neuronal subtype.

These results demonstrate that AAV-mediated gene delivery efficiency is strongly influenced by both serotype and neuronal subtype, underscoring the importance of context-specific vector selection in human neuronal models.

### Assessment of AAV-induced cytotoxicity and neurotoxicity in iPSC-derived neurons

To evaluate potential cytotoxic and neurotoxic effects of AAV transduction, we quantified both cell number and neurite outgrowth across all conditions using high-content imaging and automated image analysis (Fig. 3). Heat maps displaying total cell counts served as a proxy for general cell viability, and neurite length per cell was used as a sensitive readout of neuronal health and morphology. Analysis of the total cell counts revealed a dose-dependent reduction in viability across the most effective serotypes, suggesting that high transduction loads may induce cytotoxic effects. This negative relationship was quantified by Pearson’s correlation (R), demonstrating notable differences in cellular robustness across the derived neuronal populations.

**Figure 3:**
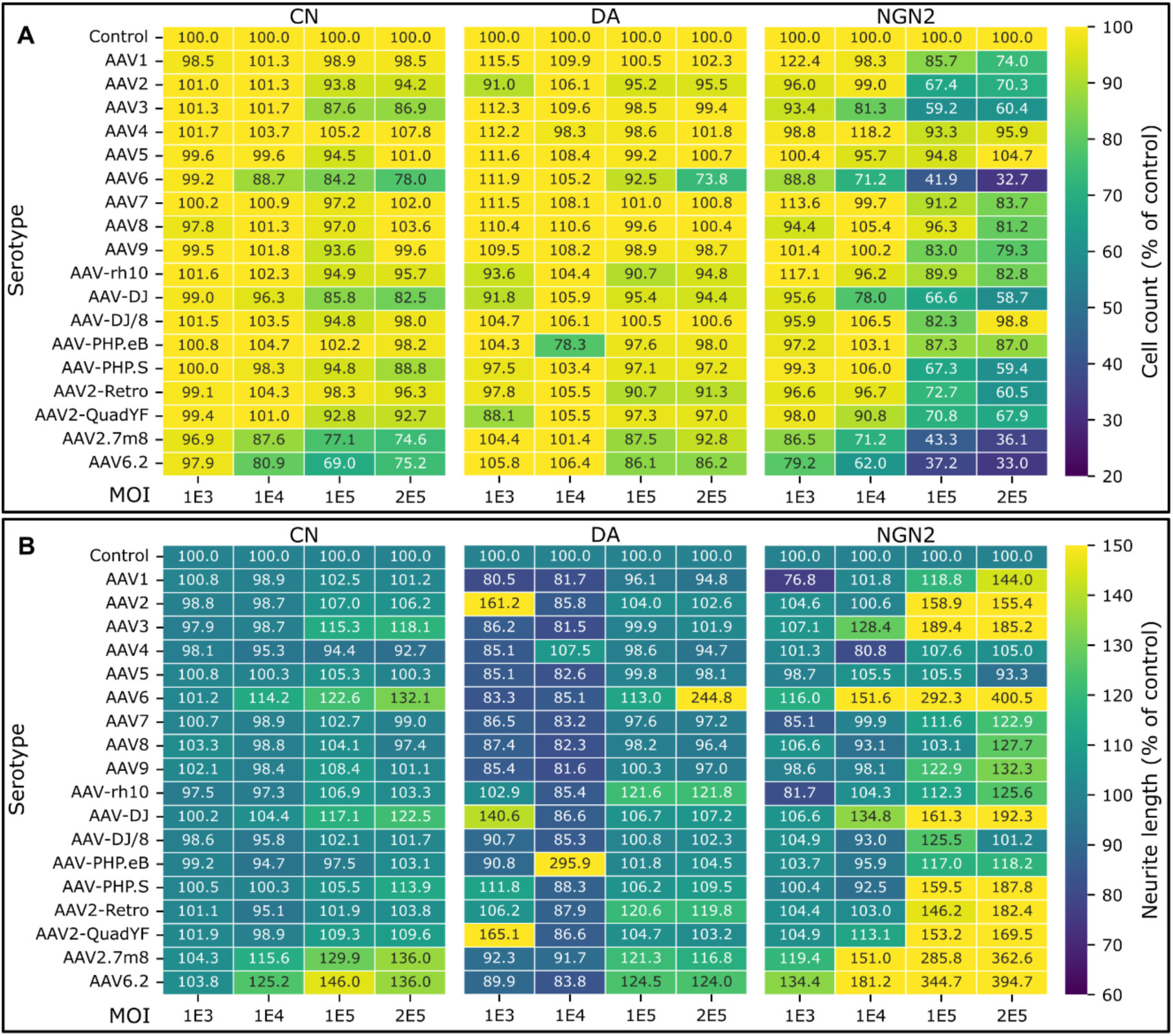
Toxicity assessment. **A:** Cell counts at endpoint determined by confocal imaging. A dose dependent decrease in cell count was observed for AAV serotypes 6, 2.7m8 and 6.2 across cell types. NGN2 neurons show stronger reduction in cell count than CN while dopaminergic neurons are the most resistant subtype. **B:** Neurite lengths per cell for all AAV dosages. Lengths were determined by confocal imaging of tubulin tracker signal. (n=3 wells, 16 FOV/well). A marked increase in neurite length per cell correlating with cell count metric can be observed.

Dopaminergic neurons (DA) were the most resilient subtype, exhibiting the weakest negative correlation between high expression and reduced cell viability. Specifically, the correlation coefficients for DA neurons were *r* = −0.36 for both GFP (%) vs. cell count and intensity vs. cell count.

Cortical neurons (CN) exhibited an intermediate level of susceptibility, as shown by stronger negative correlations: *r* = −0.86 between GFP (%) and cell count and *r* = −0.82 between intensity and cell count.

In contrast, the NGN2-derived neurons were the most susceptible to toxic side effects, displaying the strongest negative correlations between increased viral load/expression and reduced cell viability. The R values were *r* = −0.89 for GFP (%) vs. cell count and *r* = −0.91 for intensity vs. cell count. This differential tolerance highlights the critical importance of optimizing viral doses on a cell-type-specific basis, particularly for highly sensitive populations like NGN2 neurons, which showed a substantial cell number reduction of up to 67% (AAV6 and 6.2) compared to their respective non-transduced control condition at the highest MOI (Fig. 3A).

The neurite length relative to the number of cells was assessed based on the tubulin tracker signal. This measurement followed the same trends as the cell count measurement, showing the most pronounced changes at the highest doses and with the most effective serotypes. NGN2 neurons were the most susceptible, showing the greatest changes, whereas cortical neurons exhibited the fewest differences to untreated control (Fig. 3B).

### Top-performing serotypes efficiently transduce cerebellar organoids

To determine whether the top-performing AAV serotypes identified in our 2D neuronal cultures also efficiently transduce more complex human 3D models, we tested AAV2.7m8 in iPSC-derived cerebellar organoids. We analyzed both whole-mount and dissociated organoids following viral transduction at different doses (Fig. 4). Confocal imaging of cryosectioned and counterstained organoids revealed robust GFP expression up to 75% of the organoid area in whole mount organoids, with AAV2.7m8 displaying particularly strong and widespread transduction across organoid regions, although with a gradient towards the core (Fig. 4A and B). To gain deeper insights into potential cell type preferences, we used a dissociated organoid model, that retains the cellular complexity but offers improved possibilities for single-cell resolution imaging. At an MOI of 1E4 gc/cell (calculated on the seeding density), eGFP positive neural precursor cells (SOX2), neurons (MAP2), early (S100β) and late (GFAP) astrocytes could be observed. No apparent cell type preference was observed while a significant proportion of cells could not be clearly attributed to a definitive cell type by immunostaining (Fig. 4C).

**Figure 4:**
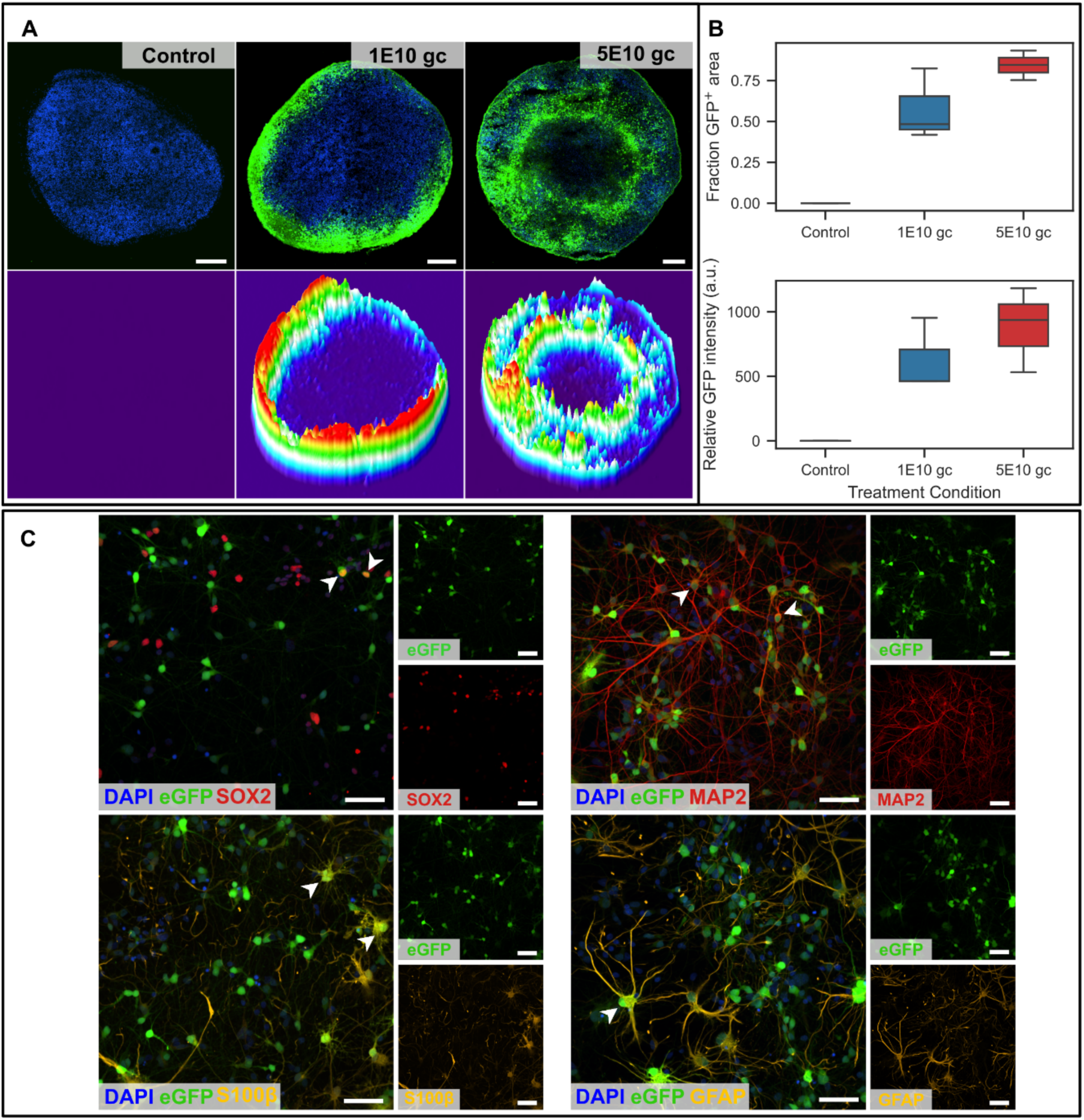
Transduction of cerebellar organoids. **A:** Whole organoid slices after transduction with 2.7m8. 3D surface plots visualize high penetrance of AAV2.7m8 as well as a transduction gradient towards the core of the organoid (Scale bar: 50 µm) **B:** Quantification of GFP+ area and signal intensity in sectioned organoids **C:** Dissociated organoid cultures after transduction with AAV 2.7m8 showing transduction of neurons, astrocytes as well as neural precursor cells (arrowheads; scale bar: 50 µm).

These findings are consistent with prior observations in related organoid systems: for example a recent study using cortical and cerebral organoids also identified AAV2.7m8 as a highly efficient serotype in 3D human neural tissue (Drouyer et al., 2024). Despite differences in organoid type and differentiation protocol, our results independently confirmed the superior performance of AAV2.7m8 in 3D brain organoids, supporting its potential as a broadly applicable vector for human CNS models.

Together, these results validate our high-throughput neuronal screen and demonstrate that the top-performing serotypes retain strong transduction capabilities in physiologically relevant 3D neural models.

## Discussion

### Context-dependent AAV performance in human neuronal subtypes

In our study, we systematically evaluated 18 AAV serotypes across four dosages in three distinct human iPSC-derived neuronal subtypes (cortical, dopaminergic and NGN2 neurons), revealing a dependence of optimal vector performance on the specific neuronal population and viral dose. We identified high-performing variants that were either effective across neuronal subtypes (AAV6, 6.2 and 2.7m8) as well as variants that were highly cell-type-specific (AAV2-retro in DA neurons). A differential susceptibility to cytotoxicity was observed when comparing the neuronal subtypes: high gene expression rates were associated with reduced viability in CN and especially NGN2 neurons (*r* ≈ −0.8 to −0.9), while DA neurons were significantly more resilient (correlation *r* = −0.36). This points towards a general GFP toxicity rather than an AAV-mediated toxicity, with NGN2 neurons being more vulnerable. In some highly toxic conditions, neurite length normalized per cell appeared increased despite reduced cell counts. One plausible explanation is survivor and density effects: if the most vulnerable neurons die, the remaining cells may represent a more resilient subpopulation with relatively greater arborization, and reduced culture density can relieve contact-dependent constraints on neurite extension, increasing neurite length per remaining cell. Finally, we successfully validated the superior transduction capabilities of one of the top-ranking serotypes, AAV2.7m8, in more complex 3D cerebellar organoids. Together, these results emphasize the critical need for context-specific vector selection and dose optimization when utilizing AAVs in human neuronal models.

### Species-specific discrepancies and unexpected tropism

The presented findings reveal crucial species- and cell-type-specific differences in AAV tropism, underscoring the necessity of using human iPSC-based models in clinical research. Serotypes that were considered ideal for the CNS in rodent models (specifically AAV9, AAV-PHP.eB, and AAV-S) showed unexpectedly poor transduction in human neurons. Conversely, some vectors not initially associated with neural tropism performed exceptionally well. For example, AAV6.2, despite being developed for lung applications (Kang et al., 2020), demonstrated exceptional and broad-spectrum transduction across our human neuronal subtypes. Additionally, AAV.DJ, tailored for transduction of the liver (Grimm et al., 2008), also performed well in iPSC-derived neurons, consistent with recent reports on its neural tropism (Chauhan et al., 2024). A more detailed investigation of AAV2.7m8 revealed superior efficiency in both 2D neuronal cultures and 3D organoids, consistent with its strong performance recently reported in other human organoid systems (Drouyer et al., 2024); suggesting a generalized high tropism for human neural tissue. Most compellingly, the highly specific performance of AAV2-retro in DA neurons, outperforming all others in expression intensity, aligns well with its selection from a library that was injected into the substantia nigra (Tervo et al., 2016). These results confirm that AAV2-retro as an optimal vector for this cell type and validate the robust identity of the DA neurons in our iPSC platform. Collectively, these observed discrepancies and unexpected positive outcomes underscore that our understanding of AAV tropism and biology remains incomplete, thus highlighting the critical importance and utility of human iPSC-derived platforms in uncovering the molecular mechanisms of species- and cell-type-specific infection.

### Translational implications for gene therapy and disease modeling

These results have direct, critical implications for *in vitro* applications as well as the development of future CNS gene therapies. Subtype-specific transduction profiles allow researchers to use the optimal vectors for disease modeling. For example, AAV2-retro specifically transduces DA neurons. Selecting and using the best vector ensures that genetic tools, such as CRISPR or optogenetics, are efficiently delivered to the targeted neuronal subtype. The validated, top-performing serotypes, particularly the broad-spectrum AAV6.2 and AAV2.7m8 and subtype-specific AAV2-retro, are now ready for application in various neural models. These vectors can be leveraged to deliver a wide range of candidate gene therapy transgenes and editing tools efficiently, allowing researchers to conduct detailed analysis of transgene subcellular localization, chronic expression profiles, and specific, long-term toxicity in patient-derived models. This is critical for developing personalized medicine approaches and tackling currently hard-to-transduce iPSC-derived neural cell types, while also providing the necessary foundation for utilizing these top serotypes to deliver diverse genetic tools in high-throughput assays, accelerating the discovery of drug targets or disease mechanisms. Crucially, our findings show that rodent-optimized “brain” serotypes (AAV9, PHP.eB, PHP.S) performed poorly in human neuronal cells implicating significant room for improvement in the development of gene therapy vectors for the CNS. Our data supports the notion that human iPSC-based CNS models have the potential to become essential tools that can effectively supplement and refine classical *in vivo* screenings, thereby streamlining the vector selection process. Moreover, the high-throughput potential inherent in iPSC-derived neuronal models allows for the rapid screening of vast AAV libraries and genetic modifications, providing an unprecedented opportunity to better understand the mechanisms of AAV infection and tropism specificity in human brain cells. The transduction profile of diverse serotypes in different neuronal subtypes provides a foundation for the rational design of next-generation vectors. For instance, combining the demonstrated superior neural cell transduction capabilities of a vector like AAV2.7m8 with the blood-brain barrier (BBB) crossing efficiency of novel serotypes (Goertsen et al., 2022) could potentially yield a therapeutic candidate superior to current vectors. This refined approach significantly increases the likelihood of success and translatability of CNS-targeted gene therapies.

### Study limitations and future perspectives

However, our study is constrained by the inherent limitations of the iPSC-derived neuronal system and our high-throughput screening methodology. The *in vitro* cultures, whether 2D or 3D, lack critical *in vivo* elements such as the blood-brain barrier (BBB), neurovascular units, and complex native immune cell interactions, preventing the assessment of vectors designed for systemic delivery (e.g., AAV-PHP.eB). Furthermore, the simplified cellular environment may not accurately replicate the expression or positioning of critical cell-surface glycoproteins or receptors found on mature primary human neurons. Methodologically, the screen was limited to a single reporter gene (GFP), meaning results may not perfectly translate when using larger therapeutic transgenes. Crucially, while our work identifies optimal serotypes for high-dose *in vitro* applications, the absolute efficiency achieved remains relatively low compared to other viral vectors, necessitating high viral loads to achieve widespread transduction. This inefficiency strongly suggests the presence of significant, unidentified cellular host restrictions within human neurons that limit AAV transduction and transgene expression. Addressing these underlying biological barriers will be necessary to unlock the full potential of AAVs for CNS disease modelling and testing of therapeutic approaches.

## Conclusion

Taken together, our systematic screen in human iPSC-derived neuronal subtypes reveals that optimal AAV performance is critically dependent on serotype, dose, and neuronal population. We identified highly effective broad-spectrum vectors (AAV6, AAV6.2, AAV2.7m8) and demonstrated subtype-specific transduction of AAV2-retro in dopaminergic neurons. Crucially, the poor performance of traditional rodent-optimized vectors highlights a major species-specific barrier. This work provides an essential, quantitative resource for *in vitro* gene delivery and confirms human iPSC-based neuronal models as valuable, high-throughput tools for species- and cell-type-specific vector selection, with the potential to accelerating the rational design and translational success of next-generation AAV gene therapies targeting the human CNS.

## Experimental procedures

### Cell lines

For all experiments presented in this study, the reference iPSC cell line KOLF2.1J was used. The KOLF2.1J line was derived from a caucasian male and was generated by and obtained from The Jackson Laboratory (Pantazis et al., 2022). Material transfer agreement between the Jackson Laboratory and the German Center for Neurodegenerative Diseases has been obtained prior to conducting any experiment.

### iPSC culture

iPSCs were cultured using established methods as previously described. In brief iPSC were maintained in essential 8 medium (E8) or Essential 8-flex medium (ThermoFisher Scientific) on growth factor reduced Matrigel (Corning) coated plated and passaged using PBS/EDTA. iPSCs were used up to passage 20.

### Cortical neurons

The protocol for the differentiation of cortical neurons was adapted from previously published articles (Rehbach et al., 2019; Shi, Kirwan, & Livesey, 2012) with minor modifications. In brief, iPSCs were seeded at high confluency and subjected to dual SMAD inhibition for 8 days followed by expansion of neural progenitors. Maturation in neuronal medium including 4 days of growth factor inhibition by DAPT and PD yielded a neuronal culture highly enriched for cortical markers after 36 days. A detailed description of the protocol was published before (Hauser et al., 2020). Neurons were used for AAV transduction experiments between day 36 and 40 of differentiation.

### NGN2 neurons

A stable smNPC line for inducible NGN2 expression and the differentiation protocol into neurons were described previously (Czuppa, Dhingra, Zhou et al., 2022). In brief, a stable smNPC line engineered for inducible NGN2 expression was generated using a single lentiviral vector; cells were used within eight passages post-transduction. The plates were coated overnight with 20 µg/mL dPGA (DendroTEK Biosciences, Montreal, Canada), washed three times with PBS, and then coated with Matrigel (1:150, Corning) for 1 h; this combined coating is hereafter referred to as dPGA/Matrigel. All coating and washing steps were performed using a MultiFlo FX dispenser (BioTek). For differentiation, cells were plated at 40,000 cells/well on PhenoPlate 96-well plates (Revvity) in N2B27 medium supplemented with 2.5 µg/mL doxycycline (to induce NGN2 expression), 2 µM DAPT and 5 µg/mL laminin, using a MultiFlo FX dispenser.

### Dopaminergic neurons

Dopaminergic neurons were generated in two steps. First, smNPCs were derived from Kolf2.1J iPSCs (https://www.jax.org/jax-mice-and-services/ipsc/) using a small-molecule protocol adapted from Reinhardt et al. (2013) and described in detail in (Dhingra et al., 2020). Second, smNPCs were differentiated into midbrain dopaminergic (mDA) neurons using an adapted small-molecule protocol based on (Reinhardt et al., 2013) as follows. smNPCs were plated on Matrigel (1:50) at 3.0 × 10^5 cells/cm^2 in N2B27 basal medium supplemented with ascorbic acid (64 µg/ml, Sigma), FGF8 (100 ng/ml, Peprotech), and purmorphamine (1 µM, Cayman). Medium was replaced on days 3 and 6 using the same patterning medium. Day 8 precursor cells were dissociated with Accutase and replated at 40,000 cells/well onto dPGA/Matrigel coated 96 well plates in N2B27 medium containing ascorbic acid (64 µg/ml), FGF8 (100 ng/ml), purmorphamine (0.5 µM), Thiazovivin (1 µM), and laminin (3 µg/ml). When necessary patterned precursors were cryopreserved on day 7 and stored long term, then thawed and reintroduced into the protocol at the replating stage (day 8) without altering subsequent medium compositions or the maturation timeline. From day 10 onward, cultures were maintained in maturation medium (N2B27 with ascorbic acid, 64 µg/ml; DAPT, 10 µM; BDNF, 10 ng/ml; GDNF, 10 ng/ml; TGF-β3, 1 ng/ml; and laminin, 1 µg/ml), with medium changes every 3 days. AAV transduction of mDA neurons was done at day 15 of differentiation.

### Cerebellar organoid differentiation

Cerebellar organoid differentiation was adapted from different previous published protocols (Muguruma et al., 2015; Nayler et al., 2021; Silva et al., 2021). Therefore, iPSCs were washed once with PBS and subsequently incubated for 8-10 min with Accutase at 37°C to get a single cell suspension. The reaction was stopped with DMEM, counted and centrifuged for 5 min at 300 *g*. Cells were resuspended in E8 + Y-27632 (20 µM, Selleckchem) and 1.8×10^6^ seeded into one 24-well AggreWell800 (pre-treated with anti-adherence rinsing solution, both StemCell Technologies). The plate was then centrifuged at 100x*g* for 1 min and subsequently placed into the incubator. The next day, medium was changed to E8 + Y-27632 (10 µM) and to growth factor chemically defined medium GfCDM + Y-27632 (10 µM) + SB431542 (Sigma, 10 µM) + LDN193189 (Th. Geyer, 333 nM) + FGF2 (Peprotech, 50 ng/ml) the following day. GfCDM) consisted of 49.5% IMDM (Gibco), 49.5% Ham’s F-12 (Gibco), 1% Chemically defined lipid concentrate (CDLC, Gibco) with BSA (5 mg/ml, Sigma), Apo-transferrin (15 µg/ml, Sigma), Insulin (7 µg/ml, Sigma), and α-thioglycerol (450 µM, Sigma). Between day 3 and 6, medium was changed to gfCDM + SB + LDN + FGF2 on a daily basis. On day 7, embryoid bodies (EBs) were transferred to a T-75 (Corning) flask, pre-treated with anti-adherence rinsing solution. From now on EBs were incubated on an orbital shaker (50 rpm) at 37°C. Medium was changed every other day to gfCDM + SB431542 (6.76 µM, Sigma) + LDN193189 (222 nM, Th. Geyer) + FGF2 (33.3 ng/ml) between day 7 and 13, and gfCDM + FGF19 (100 ng/ml, Peprotech,) until day 20. Between day 21 to 34, medium was changed three times a week to Neurobasal-based medium (NBM) consisting of Neurobasal (Gibco) + 1% N2 (Gibco) + 1% Glutamax (Gibco) + 1% PenStrep (Gibco) + 0.1% AmphoB (Gibco). For the second week of NBM, medium was supplemented with SDF-1 (300 ng/ml, Peprotech). Between day 32 and 61, medium contained 0.66% matrigel (Corning). From day 35 on, medium was changed to BrainPhys with 2% SM1 (both StemCell Technologies) + 1% N2 (Gibco) + 1% PenStrep (Gibco) + 0.1% AmphoB (Gibco) + BDNF (20 ng/ml, Peprotech) + GDNF (20 ng/ml, Peprotech) + dbcAMP (1 mM, Sigma) + Ascorbic acid (Tocris, 200 µM).

### Dissociation of cerebellar organoids

Cerebellar organoids were dissociated using the GentleMACS neural tissue dissociation kit (Milteneyi Biotec) according to manufacturer’s instructions. Single cell dissociation was ensured using a 70 µm cell strainer (Milteneyi Biotec) and cells were seeded in a density of 150000 cells/cm^2^ to Poly-L-ornithine/Matrigel (1:30) coated tissue culture plates. Cells were allowed to mature for 14 days in organoid medium before AAV treatment, fixation, immunostaining and analysis by confocal imaging.

### AAV transduction

A panel of 18 AAV serotypes harboring identical CAG-eGFP expression constructs (VB010000- 9287ffw) was purchased from vectorbuilder (PANEL-AAVS02-41). All serotypes were diluted to an equal titer of 1E13 gc/ml in the virus buffer (PBS + 200 mM NaCl + 0.001% Pluronic F- 68). The master AAV plate was prepared by diluting in 3N neuronal medium to a concentration of 1E13 gc/ml in a 96 well U bottom plate. From this plate, the dilutions for the four tested MOIs were prepared using a 96 well pipettor head (Integra) and subsequently added to the prepared 96 well (PhenoPlate 96-well black, 6055308, Revvity) imaging plates. Medium was changed 48h later using MultiFlo FX dispenser (BioTek). The highest dosage was monitored for GFP expression using an Incucyte microscope system (Sartorius) acquiring images every 6 hours after medium change. For confocal screening all cells were allowed to express GFP until day 5 after transduction. Cerebellar organoids were transduced as described in (Drouyer et al., 2024).

### qPCR

Mature neurons were harvested and RNA was isolated using GeneJet RNA isolation kit (ThermoFisher Scientific) according to manufacturer’s instructions with the addition of on column DNA digestion step (Qiagen RNAse free DNAse Kit). 500 ng of RNA was reverse transcribed using Revert Aid first strand cDNA synthesis Kit (ThermoFisher Scientific) and used for qPCR assay. For qPCR, SYBR Select Master mix (Invitrogen) was used together with respective primer pairs in the ViiA7 qPCR system. List of primers used for qPCR can be found in supplemental table S3.

### Live-cell staining

96-well plates were incubated with a staining solution containing TubulinTracker FR (0.125 µM, ThermoFisher Scientific) and Hoechst 33342 (1 µg/ml, ThermoFisher Scientific) for 1 h at 37°C in 3N neuronal medium, followed by a medium change to 3N neuronal medium. Cells were allowed to equilibrate at 37°C for 3 h min before automated imaging.

### Immunostaining

Cells were fixed using 4% paraformaldehyde for 10 min at 37°C, permeabilized and blocked with 5% bovine serum albumin (BSA) in PBS containing 0.1% Triton X-100 (PBS-T). Primary antibodies were diluted in the same blocking solution and incubated overnight at 4°C. After washing steps, matching secondary antibodies were applied in PBS-T in a dilution of 1:1000 for 45 min at RT followed by 15 min of DAPI/RNAseA treatment for counterstaining. For imaging, wells of 96-well plates were filled with PBS. Cover glasses were mounted using DAKO mounting medium and allowed to dry overnight. Antibodies used for immunostaining are listed in supplemental table 1.

Samples were imaged using the Nikon Ti2e-SORA system equipped with the Yokogawa W1 spinning disc unit and 40x water immersion objective. Ten fields were selected manually and imaged in 25 µm z-stacks with subsequent maximum intensity projection.

### Confocal imaging

The neurons in 96 well plates were subjected to automated imaging using CV7000 (Yokogawa) after live-cell staining protocol from above. The images were acquired using predetermined settings using 20x objective for positive controls and untransduced cells (Dhingra et al., 2020) Per condition, n = 3 wells were imaged with 16 fields of view per well using fixed acquisition settings across plates and serotypes.

## Data analysis

Confocal live cell data generated by the CV7000 imaging platform for screening of the 18 serotype AAV panel was analyzed by Cell pathfinder software (Version: 3.07.02.02, Yokogawa) to quantify GFP-positive cells, GFP intensity, total cell counts, and neurite length per cell. Live cell imaging data was processed using the Incucyte 2023A Rev2 software package (Sartorius). The resulting tables were further processed and plotted using the python libraries numpy, seaborn and matplotlib. Pearson correlation coefficients were used to assess associations between transduction/expression metrics and toxicity readouts across conditions.

## Author contribution

The study has been designed by LW, AD and SH. Differentiation of cortical neurons and cerebellar organoids has been performed by LW with the help of LR and JH. Differentiation of NGN2 and dopaminergic neurons has been performed by AD, SRN and AL. LW and AD performed the AAV serotype testing including data analysis. Gene expression analysis and ICC has been performed by LW. The manuscript has been prepared by LW, AD and SH. All authors have reviewed and approved the manuscript.

Conceptualization: LW, AD, SH

Methodology: LW, AD, SRN, AL, LR, JH

Formal Analysis: LW, AD, SH

Investigation: LW, AD, SRN, AL, LR, JH

Data Curation: AD, LW

Writing – Original Draft: LW

Writing – Review & Editing: AD, SH

Visualization: LW, AD

Supervision: LS, TG, AD, SH

Project Administration: SH

Funding Acquisition: LS, TG, AD, SH

## Acknowledgments

This study has been supported by the German Research Foundation (DFG, grants SCHO754/8- 1, HA9848/2-1) and the DZNE Stiftung – Forschung für ein Leben ohne Demenz, Parkinson und ALS (to AD, https://www.dzne-stiftung.de/).

