## Supplementary Files for "Comprehensive investigation of AAV tropism across human iPSC-derived neuronal subtypes"

### Supplementary Figures

### Figure S1:

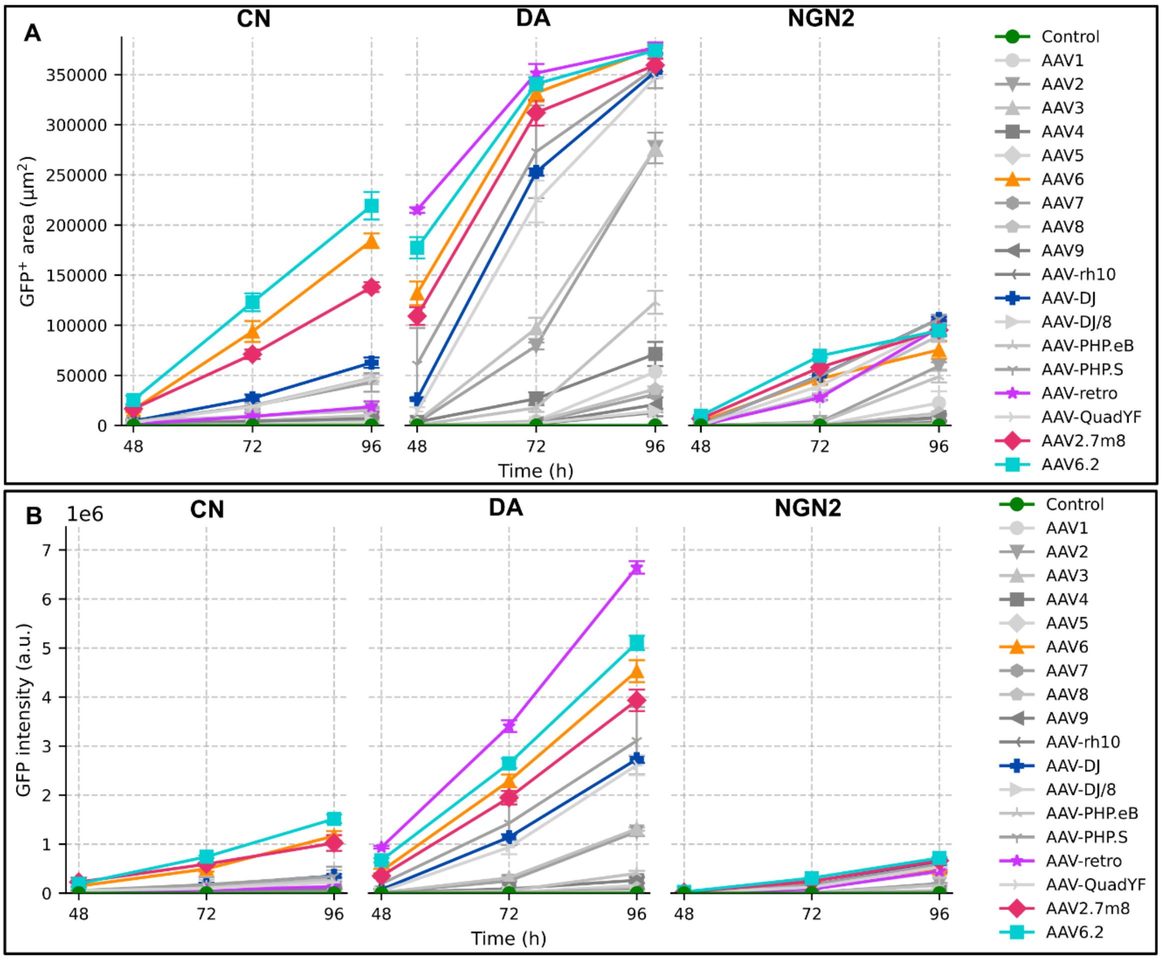


***Figure S1: Live cell imaging of highest dose (MOI 2E5). A:*** *GFP+ area over time for cortical (CN) dopaminergic (DA) and NGN2 neurons. Best performing serotypes are highlighted.* ***B:*** *GFP intensity (a.u.) over time, best performing serotypes are highlighted.*

**Figure S2:**

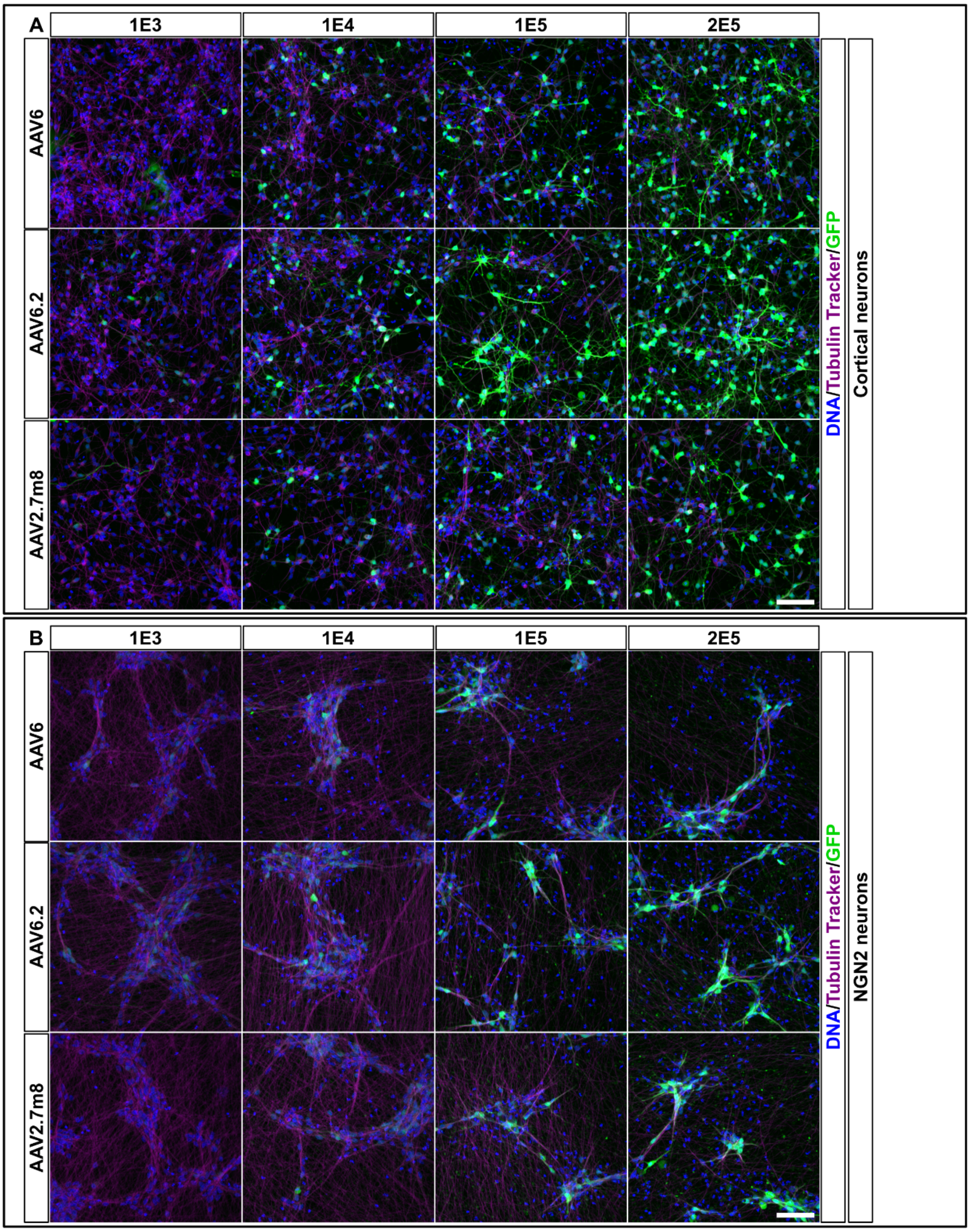


***Figure S2: Exemplary confocal images for selected serotypes in different AAV dosages (MOIs)**: **A:** Cortical neurons and **B:** NGN2 neurons transduced with AAV6, 6.2 or 2.7m8, respectively. Cells were labelled with the live cell dye Tubulin Tracker Deep red (purple) and counterstained with Hoechst (DNA, blue) before imaging (Scale bar = 50 µm).*

**Figure S3:**

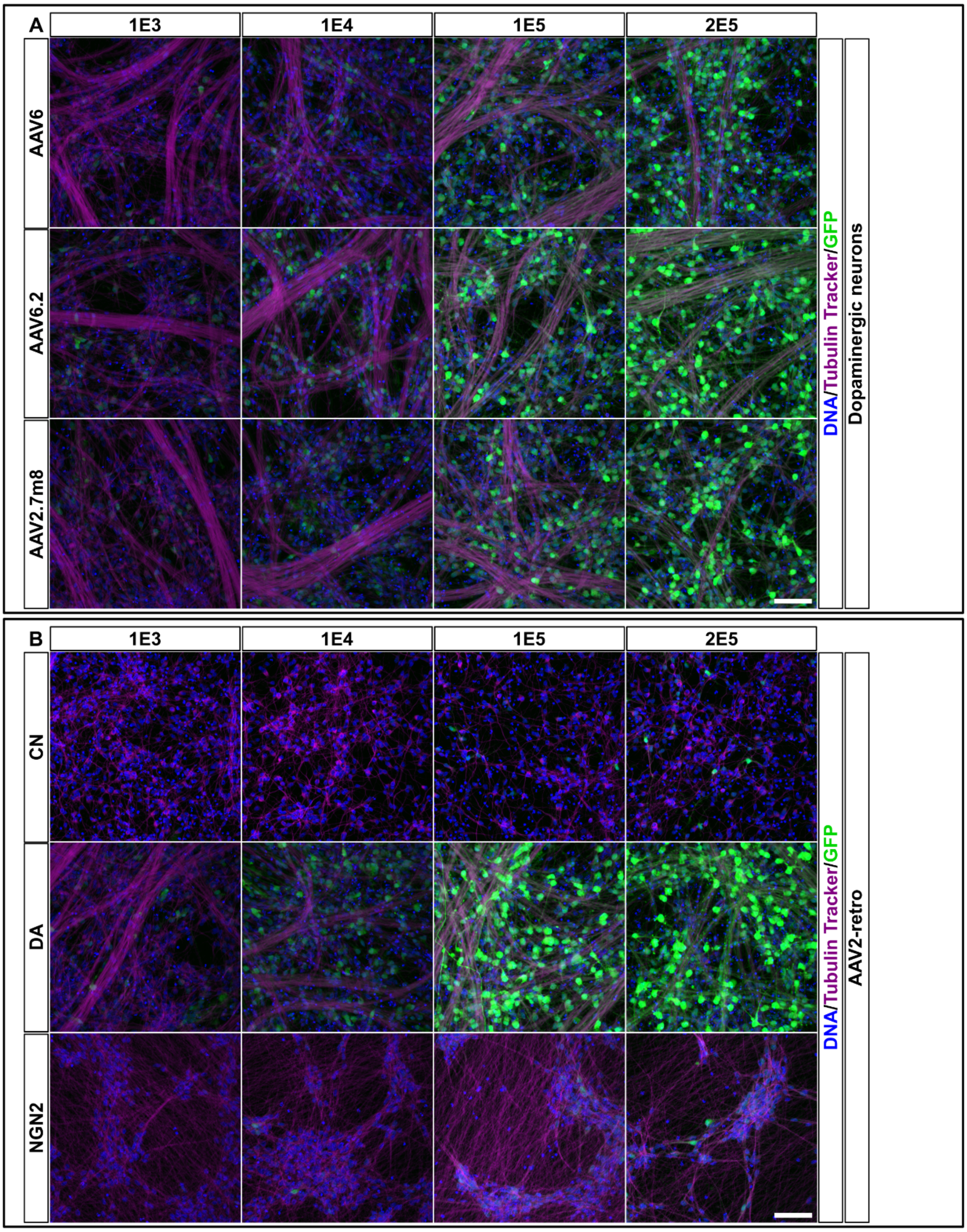


***Figure S3: Exemplary confocal images for selected serotypes in different AAV dosages (MOIs): A:*** *Dopaminergic neurons transduced with AAV6, 6.2 or 2.7m8 respectively.* ***B:*** *AAV2-Retro transduced Cortical, DA and NGN2 neurons. Specifically, DA neurons show high reporter gene expression. Cells were labelled with the live cell dye Tubulin Tracker Deep red (purple) and counterstained with Hoechst (DNA, blue) before imaging (Scale bar = 50 µm).*

### Supplementary Tables

#### Primary Antibodies

*Table S1 List of primary antibodies used in this study.*

| Target | Supplier | Cat. # | Dilution |
|---|---|---|---|
| TUJ | Sigma | T8660 | 1:1 000 |
| MAP2 | Biolegend | 822501 | 1:5 000 |
| CTIP2 | Abcam | ab18465 | 1:500 |
| TH | CellSignaling | 58844 | 1:200 |

#### Secondary antibodies

*Table S2 List of secondary antibodies used in this study*

| Name | Reactivity | Fluorophore | Suplier | Cat # |
|---|---|---|---|---|
| Goat anti-rabbit IgG (H+L) | rabbit | Alexa488 | Invitrogen | A-11008 |
| Goat anti-rat IgG (H+L) | rat | Alexa568 | Invitrogen | A-11077 |
| Goat anti-Mouse IgG (H+L) | mouse | Alexa647 | Invitrogen | A-21235 |
| Goat anti-chicken IgG (H+L) | chicken | Alexa647 | Invitrogen | A-21449 |

#### qPCR Primers

*Table S3:List of qPCR primer sequences used in this study*

| Target | Forward Primer (5’->3’) | Reverse Primer (5’->3’) |
|---|---|---|
| TUJ | TGGACATCTCTTCAGGCCTG | ATGATGCGGTCGGGATACTC |
| TAU | CCTCTGATCCAACCCTCCAG | TTTTGCTGGAATCCTGGTGG |
| BRN2 | ACGACGAGCCACACCATG | CTTGAACTGCTTGGCGAACT |
| NESTIN | TGCGGGCTACTGAAAAGTTC | GGCTGAGGGACATCTTGAG |
| CTIP2 | GGTCTGGAGATAGAGGAGCC | TGTCCAGGGCCTTGTCATAG |
| SYN | CGATGCCAAATATGACGTGCGTG | AGCATCGCAGAGCCAGTATTGG |
| TH | CCTGTACTGGAAGGCGATCTCA | GCTGGACAAGTGTCATCACCTG |
| GIRK | GAAGGAACTCACCCTCAGGTGT | GATCTCCATGAGGGACGGAAAAC |
